# Myelodysplastic syndromes disable support of human hematopoietic stem and progenitor cells by CD271, VCAM- 1 and CD146 expressing marrow niche cells

**DOI:** 10.1101/2023.04.09.536176

**Authors:** Yuko Kawano, Hiroki Kawano, Dalia Ghoneim, Thomas J. Fountaine, Daniel K Byun, Mark W. LaMere, Jason H. Mendler, Tzu-Chieh Ho, Noah A. Salama, Benjamin J. Frisch, Jason R. Myers, Siba El Hussein, John M. Ashton, Mitra Azadniv, Jane L. Liesveld, Youmna Kfoury, David T. Scadden, Michael W. Becker, Laura M. Calvi

**Author notes:** equal contributed author (Yuko Kawano, Hiroki Kawano).

## Abstract

Mesenchymal stem/stromal cells (MSCs) within the bone marrow microenvironment (BMME) support normal hematopoietic stem and progenitor cells (HSPCs). However, the heterogeneity of human MSCs has limited the understanding of their contribution to clonal dynamics and evolution to myelodysplastic syndromes (MDS). We combined three MSC cell surface markers, CD271, VCAM-1 (Vascular Cell Adhesion Molecule-1) and CD146, to isolate distinct subsets of human MSCs from bone marrow aspirates of healthy controls (Control BM). Based on transcriptional and functional analysis, CD271^+^CD106^+^CD146^+^ (NGFR+/VCAM1+/MCAM+/Lin-; **NVML**) cells display stem cell characteristics, are compatible with murine BM-derived Leptin receptor positive MSCs and provide superior support for normal HSPCs. MSC subsets from 17 patients with MDS demonstrated shared transcriptional changes in spite of mutational heterogeneity in the MDS clones, with loss of preferential support of normal HSPCs by MDS-derived NVML cells. Our data provide a new approach to dissect microenvironment-dependent mechanisms regulating clonal dynamics and progression to MDS.

## INTRODUCTION

The BM microenvironment (BMME) serves as the milieu that maintains normal hematopoiesis via cell-cell interactions and secretory factors^1^. Non-hematopoietic stromal cells including mesenchymal stem/stromal cells (MSCs), osteolineage cells (OLC), and bone marrow (BM)-derived endothelial and perivascular cells contribute to regulation of hematopoietic stem cell (HSC) function^2–5^. Dysfunction in some of these populations has been reported to participate in the development of hematological malignancies including myelodysplastic syndrome (MDS) and leukemia^6–9^. We have previously shown in a murine model of MDS, that targeting the BMME can mitigate MDS associated marrow failure and disease progression, decreasing and delaying the risk of transformation to leukemia^10^. MSCs are a key HSC niche cell population in the BMME. The definition of MSCs is evolving. MSCs currently are thought to be a heterogeneous population of non-blood forming stem and progenitor cells with potential to differentiate into multiple cell lineages, including osteoblastic, chondrocytic, and adipocytic cells^11^. Although prior studies and consensus statements have reported surface markers for isolating MSCs^12,13,14^, recent findings demonstrated significant diversity in murine BM stroma populations utilizing single cell RNA-seq (scRNA-seq)^15^ not accounted for in the study of human BMME populations. While prospective isolation of murine MSC relies on positive expression of CD51 (Integrin αV)^16^ and Sca1 (stem cell antigen-1)^17^, cell surface markers available for human MSC that individually enrich for stemness and for hematopoietic stem and progenitor cell support include CD271/NGFR (Nerve Growth Factor Receptor)^18^, CD146 (Melanoma Cell Adhesion Molecule)^19, 20^, and CD106 (VCAM-1)^21^. Using these markers in combination, we identified two novel subpopulations of MSCs from human bone marrow. CD271^+^CD106^+^CD146^+^ (NGFR+/VCAM1+/MCAM+/Lin-; **NVML**) cells, compared to CD271^+^CD146^-^ (NGFR+/Lineage- cells; **NLCs**). Transcriptionally and functionally, NVML cells display stem cell characteristics and provide superior support for normal HSPCs. Moreover, NVML cells from a cohort of MDS patients demonstrated conserved loss of HSPC support capacity in spite of clinical and mutational heterogeneity.

## RESULTS

### Distinctive transcriptional profiles of novel human bone marrow-derived MSC subsets

After obtaining bone marrow aspirates from eight healthy individual, we used our novel sorting platform to prospectively isolate three human BMME stromal cell populations from CD45-/CD34-/CD31-/CD235ab- cells: CD271^+^CD146^-^CD106^-^ (NLCs), CD271^+^CD146^+^CD106^-^ (NML cells), and CD271^+^CD146^+^CD106^+^ (NVML cells) (Figure 1A). Principal component analysis (PCA) of the transcriptomes from each sorted cell population demonstrated that NLCs and NVML cells are clearly distinguishable (Figure 1B), while the transcriptional profile of NML cells overlaps with the two other populations (Supplemental Figure 1A). Similarly, heatmap and hierarchical clustering analyses of differentially expressed genes showed clear separation of the transcriptomes of NLCs and NVML cells (Figure 1C, Supplemental Figure 1B). Together, these data suggest that NVML cells and NLCs represent distinct populations, while NML cells may represent a transitioning population between these two.

**Figure 1.**
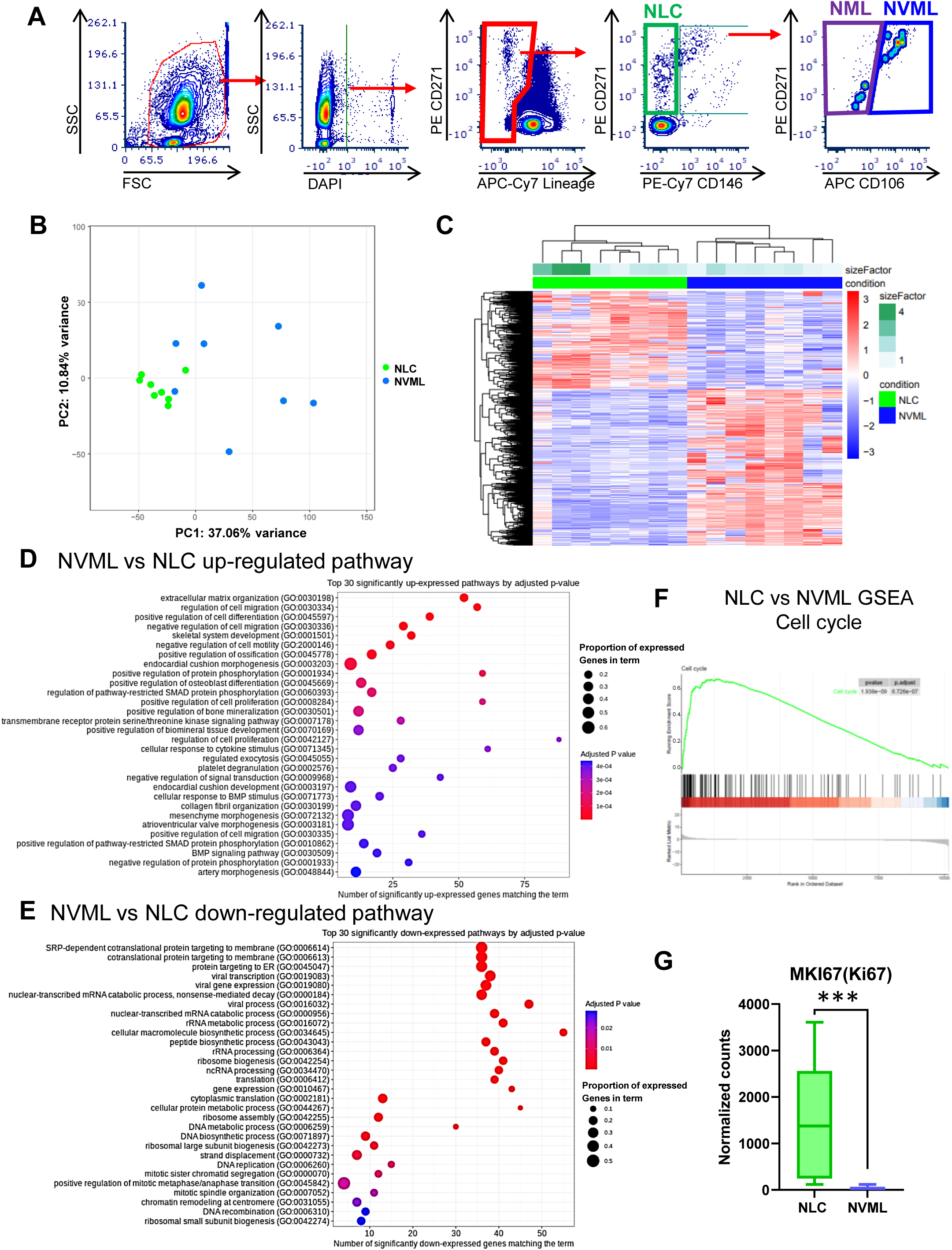
NVML cells have distinctive transcriptional profiles compared to NLCs. A. Gating strategy for sorting of NLCs and NVML cells population. B. Plot of first (PC1) and second (PC2) principal components from principal component analysis of RNA expression in Control BM samples. Each dot represents an individual patient. NLCs (green) and NVML (blue) samples. C. Hierarchical clustering heat map of differentially expressed genes in NVML cells vs NLCs samples scaled per gene. D. Up-regulated and E. down-regulated gene enrichment results from differentially expressed genes in NVML vs NLC samples. Only the top 30 hits are shown. Circle size represents proportion of differentially expressed genes enriched for that term. F. Enrichment curve of Cell cycle pathway obtained by GSEA analysis in comparison of Control NLC vs NVML transcripts. G. Normalized counts data of *MKI67* (Ki67) of NVML cells and NLCs obtained from RNA seq data analysis. n=8, Min to max, adjusted p(q) is provided. ***: q<0.001

To characterize these novel human stromal cell populations, we performed gene ontology analysis, and found that, in NVML cells, pathways associated with stemness and multiple lineage differentiation were up-regulated, while pathways related to DNA/RNA processing were down-regulated (Figure 1D, 1E). Furthermore, expression of *MKI67*, the gene encoding Ki67, a known proliferation marker, was higher with NLCs (Figure 1F). This suggests that NVML cells are more quiescent and multipotent in comparison to NLCs.

Recently the Scadden laboratory employed an unbiased approach to define the cellular taxonomy of murine non-hematopoietic BM niche cells. They reported the transcriptional profile of Leptin receptor (LepR)+ MSCs, an MSC subpopulation critical for HSC and progenitor cell support in mice.^15^ Compared to NLCs, NVML cells more closely recapitulated the LepR+ MSC subset identified in the Baryawno paper as shown by expression of key genes including *CXCL12 (SDF-1a)*, KITLG (*Kit ligand*) and *LEPR* (Figure 2A). As expected, the expression of *VCAM1,* the gene that encodes CD106, was increased in NVML cells (Fig. 2A,B). NMVL cells are also likely enriched for human CXCL12 Abundant Reticular (CAR) cells, previously shown to be a critical HSC-supportive population, since NVML have increased expression not only of *CXCL12* but also of the characteristic CAR cell genes *FOXC1* and *EBF3*^22–, 24^(Fig.2A,B). Again suggesting that NML cells represent a transitional stromal cell population, NML cells were distributed randomly amongst other populations in the heatmap (Supplemental Figure 2A). Together, these data suggest that NVML cells are enriched for cells that have the potential to support hematopoietic stem and progenitor cells.

**Figure. 2.**
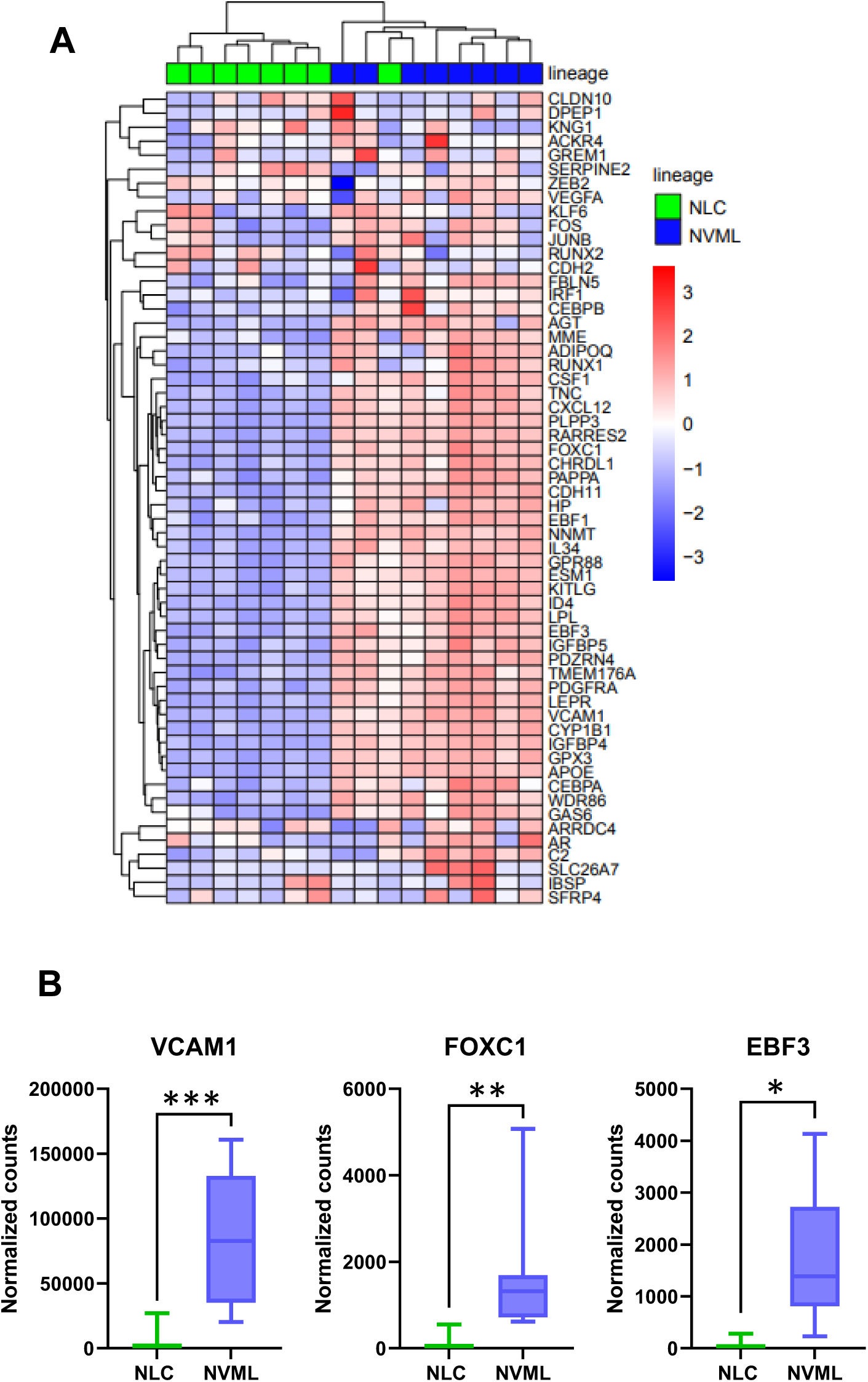
Transcriptional profile of NVML cells is compatible with murine LepR+ MSCs. A. Hierarchical clustering heat map of the set of differentially expressed genes from LepR+ MSCs in scRNAseq murine dataset^15^ in NVML cells (Blue) vs NLCs (Green) samples scaled per gene. B. Normalized counts data of key transcriptomes represent the characteristics of LepR+ MSC or CAR cells, *VCAM1, FOXC1, EBF3* in comparison of NVML cells and NLCs obtained from RNA seq data analysis. n=8, Min to max, adjusted p(q) was provided. *: q<0.05; **: q<0.01; ***: q<0.001.

### NVML cells are enriched for multipotent stem cells with trilineage differentiation capacity

We next investigated the trilineage differentiation capacity of NVML and NLC populations using *ex vivo* culture. Within the stromal cell pool, NVML cells are relatively rare (Figure 3A), however they have higher CFU-F capacity compared to NLCs (Figure 3B). Transcriptional analysis found that key markers for tri-lineage differentiation potential (osteogenic marker: *SPP1* (Osteopontin), adipogenic marker: *PPARG*, and chondrogenic marker: *SOX9*) were increased in NVML cells compared to NLCs (Figure 3C). Consistent with this, NVML cells had higher propensity for osteogenic, adipogenic, and chondrogenic differentiation compared to NLCs (Figure 3D-G). These characteristics are compatible with increased multipotentiality and stemness of NVML cells as compared with NLCs.

**Figure. 3.**
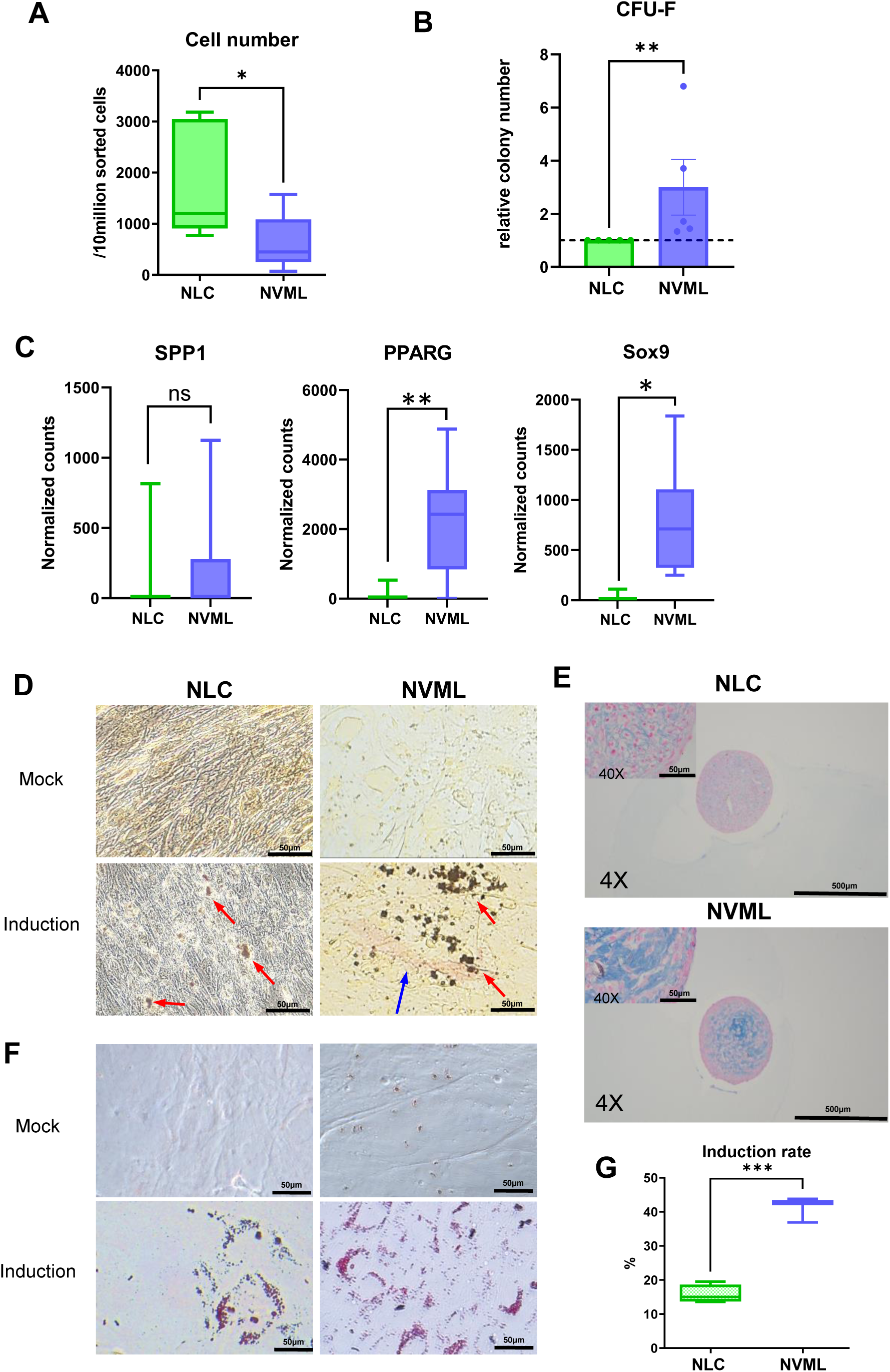
NVML cells show stem cell like characteristics compared to NLCs. A. Sorted cell number per 10 million loading (after negative selection) B. Relative CFU-F number, each dot represents an individual experiment. C. Normalized counts from RNA-seq analyses for tri-lineage differentiation gene markers (osteogenic: osteopontin (*SPP1*), adipogenic: PPAR-γ (*PPARG*), chondrogenic: SOX-9 (*SOX9*) for NVML cells compared to NLCs. D. Representative images of alkaline phosphatase/Von Kossa double staining after osteogenic differentiation of NLCs and NVML cells. Red arrows are pointing mineralizing deposits (black dots) and blue arrow is pointing to alkaline phosphatase positive area (pink region). Mock: normal media not including differentiation factors. Induction: media with differentiation factors (see methods). E. Alcian blue staining of chondrogenic differentiation test. F. Representative images of Oil-Red-O staining (red) of adipogenic differentiation test. G. Quantification of adipogenic induction rate (% of Oil-Red-O positive cells). n=3-5, Student t-test, mean and SEM or Min to max are provided in A, B and G.*: p<0.05; **: p<0.01; ***: p<0.001. Min to max, adjusted p(q) was provided in C: ns: not significant; *: q<0.05; **:q<0.01.

### NVML cells and NLCs differentially support hematopoietic stem and progenitor cells

A key function of MSC niche cells is support of normal adult hematopoiesis. In murine BMME studies, genetic tools and single cell datasets have identified subsets of BM MSCs that support hematopoiesis.^2–5, 15, 25^ Our transcriptional analysis suggest that NVML cells recapitulate the LepR+ MSC subset compared to NLCs^15^. To functionally evaluate the supporting capacity of NVML and NLC populations, we established a growth factor-free co-culture system using each sorted stromal cell subset with normal bone marrow-derived hematopoietic cells (Control BM) (Figure 4A). After 3 days of co-culture, we found that NVML cells provided superior support to Control BM CFU-Cs compared to Control BM cultured alone or with NLCs (Figure 4B). This increased ability of NVML to support HSPCs is consistent with their increased expression of the LepR and of the HSC-supportive ligands *CXCL12 (SDF-1a)*, KITLG (*Kit ligand*) and ANGPT-1 (*Angiopoietin-1*) (Figure 4C). NVMLs also demonstrate increased expression of cytokines support hematopoietic progenitors (*CXCL10*, *CCL2*, *CSF1*) (Fig 4C). These data indicate that NVML cells are niche cells for normal HSPCs.

**Figure. 4.**
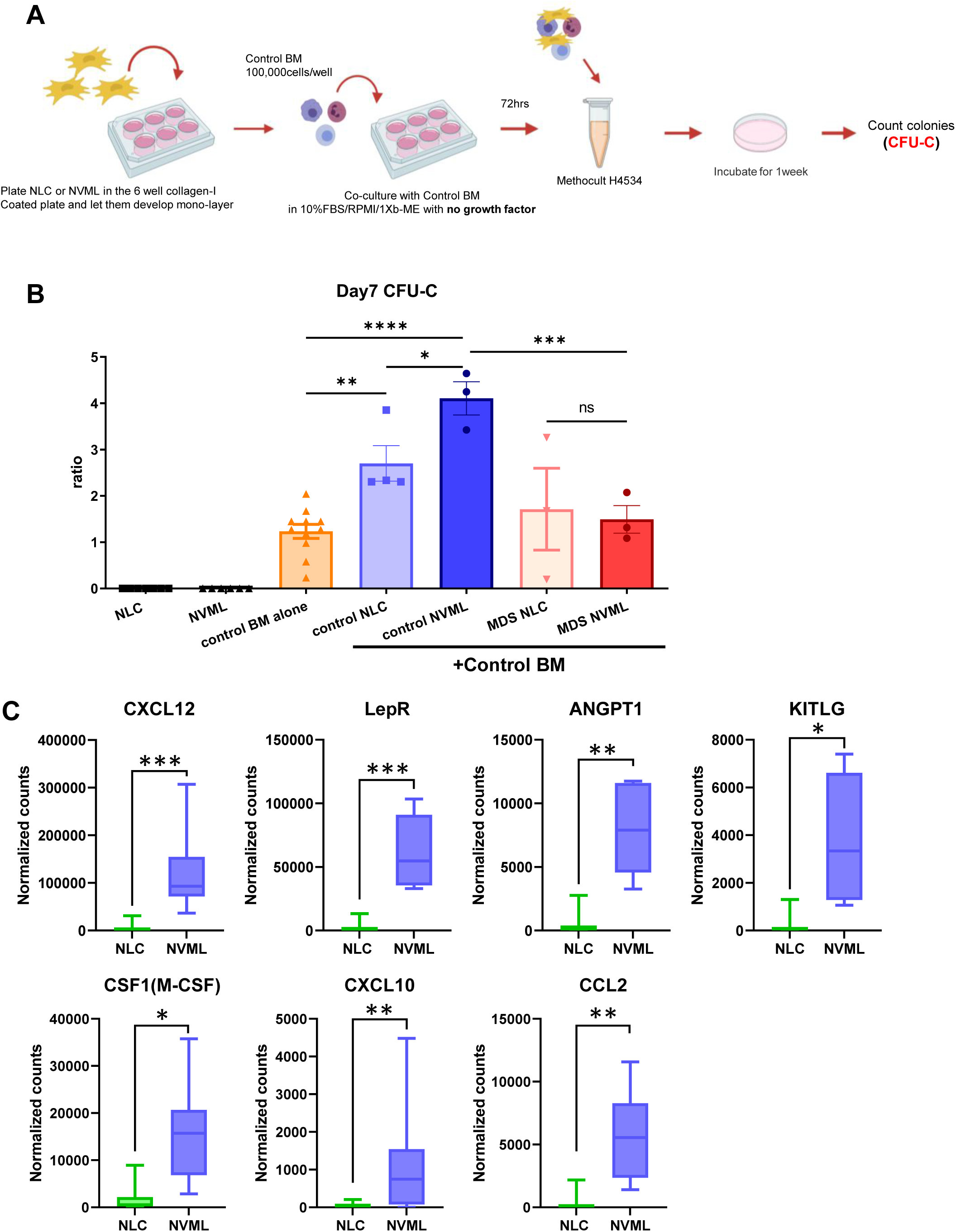
NVML cells preferentially support normal hematopoietic stem and progenitor cells. A. Schematic representation of experimental procedure of *ex vivo* co-culture of MSCs and hematopoietic cells to evaluate CFU-Cs. B. Normalized count of CFU-Cs normal NLC and NVML alone, Control BM alone (n=11), compared with Control BM supported by NVML cells or NLCs derived from Control BM (control NLC/control NVML) or MDS (MDS NLC/MDS NVML). n=3-4 separate BM-derived MSCs from Control BM or MDS patients. One-way ANOVA with corrected method of Benjamini and Yekutieli, mean and SEM.*: p<0.05; **: p<0.01; ***: p<0.001 C. Normalized counts of key transcriptional markers for hematopoietic support. Min to max, adjusted p(q) are provided. *: q<0.05; **: q<0.01; ***: q<0.001.

### NVML cells are selectively disrupted in the MDS bone marrow microenvironment

To characterize the impact of MDS on the NVML and NLC sub-populations of BM MSCs, we analyzed the transcriptome of NVML cells and NLCs from patients with MDS in comparison with aged-matched Controls. Our MDS patient cohort was clinically and mutationally heterogenous (Supplemental Table 2, Supplemental Table 3 and Supplemental Figure 3). The transcriptome of NVML cells from MDS patients was more severely disrupted compared to that of NLCs (Figure 5A). Notably, in patients with MDS, the number of differentially expressed genes (DEGs) between NVML cells and NLCs was decreased compared to Control, especially in patients clinically defined as high-risk (MDS-EB) based on the blast percentage of BM or peripheral blood (Figure 5B). Therefore, in MDS, these two microenvironmental populations become progressively more similar to each other transcriptionally in spite of being phenotypically distinct. Top hits up-regulated pathways in NVML cells in MDS compared to Controls included DNA/RNA processing and cell cycle (Supplemental Figure 4A, Figure 5C), exemplified by the upregulation of Ki67 expression in NVML cells in MDS compared to Control (Figure 5D). These data suggest that NVML from MDS patients lose stemness characteristics. Moreover, MDS-induced changes in the transcriptional profile include evidence of response to increased cellular stress and DNA damage, with enrichment in cellular senescence and Fanconi anemia pathways in NVML from MDS compared to NVML from Controls (Figure 5C, Supplemental Figure 4C)^26^. Notably, increased senescence has been previously reported in MSCs from MDS patients^6, 27, 28^. Other significantly upregulated genes in NVML in MDS were related to inflammation and metabolism (Supplemental Figure 4D). Most of the pathways downregulated in NVML cells from MDS patients were related to cytokine signaling (Supplemental Figure 4B). Importantly, the expression of some of the key HSC-supportive genes such as *CXCL12*, *KITLG* (Kit ligand), *ANGPT-1* (Angiopoietin-1), and others were downregulated in NVML cells from MDS patients (Figure 5E, Supplemental Figure 4E). Collectively, these findings would predict that, in spite of mutational heterogeneity in the MDS cells, NVML cells from MDS patients would have decreased capacity to support normal hematopoietic stem and progenitor cells.

**Figure. 5.**
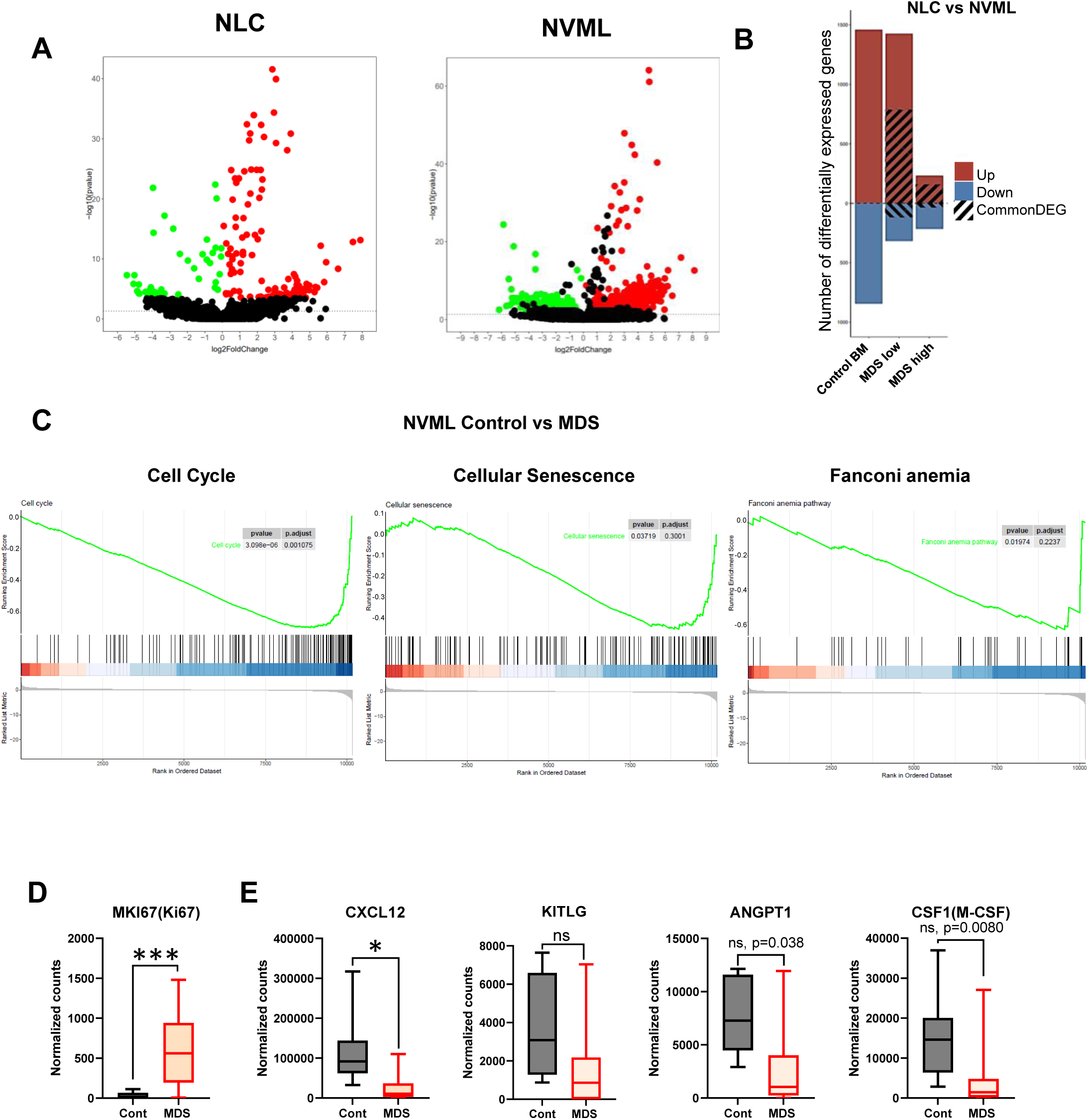
NVML cells from MDS patients are uniquely disrupted transcriptionally and functionally. A. Volcano plots based on RNAseq expression of NLCs and NVML cells from MDS compared to Control bone marrow samples. The dashed line marks p-value threshold of 0.05. Significantly up and down-regulated genes with adjusted p-value <= 0.05 are highlighted in red and green respectively. Adjusted p-value was calculated using benjamini-Hochburg correction method. B. Bar plot of number of differentially expressed genes (DEGs) between NLCs and NVML cells. Red and blue bars indicate the total number of up and down regulated genes, respectively. The striped bars in the MDS low-risk NLC vs NVML and MDS high-risk NLC vs NVML indicated the number of DEGs that overlap with Control NLC vs NVML DEGs. C. Enrichment curve of Cell cycle, Cellular senescence and Fanconi anemia pathways obtained by GSEA analysis in comparison of NVML Control vs MDS. D,E. Normalized counts for transcriptional expression of *MKI67 CXCL12*, *KITLG*, *ANGPT1*, *M-CSF* in Control (Cont) NVML cells vs MDS NVML cells. n=8-10 Min to max, adjusted p(q) or p value from Wald test (p) were provided. *: q<0.05; ***: q<0.001.

Consistent with the transcriptional analysis, coculture of MDS-derived NVML cells with Control BM demonstrated a 4 fold loss of hematopoietic stem and progenitor support compared to normal NVML (Figure 4B). In fact, NVML from MDS patients did not provide additional HSPC support compared to bone marrow alone, demonstrating MDS-dependent disabling of the HSPC niche. Moreover, compared to NLCs, NVML cells from MDS patients were no longer superior in their support of normal CFU-Cs (Figure 4C). These data suggest that a shared characteristic in the MDS microenvironment, regardless of mutation type, is loss of NVML cell function as supportive niches for normal HSPCs.

## DISCUSSION

MSCs are a critical cell component of the BMME and their roles in supporting normal hematopoiesis as well as in the pathogenesis of myeloid malignancies have been explored mainly using murine models. Recently, human bone marrow MSCs have emerged as therapeutic tools for a number of diverse applications, including immune remodeling and tissue regeneration, but whether specific MSC-dependent signals contribute to disease progression in myeloid neoplasms remains elusive. An improved understanding of the diversity of BM MSC subpopulations would significantly aid our ability to define and modulate the role of the BMME in normal and malignant hematopoiesis.

Dissecting the heterogeneity among bone and marrow stromal progenitors has been complicated by the rarity of these populations and limited availability of MSC- specific surface markers. Single cell sequencing analysis provides a promising tool for dissecting this population, but with a rarity of <1%, getting enough cells and deep enough reads to distinguish this heterogeneity is near the limit of detection. While this problem remains in murine models, CD271 (NGFR) has more recently been identified in humans as specific to a broad population of bone marrow MSCs. CD271 is a neurotrophic factor receptor known to play a role in neuron development, neuron networking^29^ and the repair of skeletal injury^30^. While expression levels on MSCs vary by report and tissue of origin, CD271 is highly expressed on human MSCs isolated by *ex vivo* culture of plastic-adherent cells and on BM-derived MSCs^31^.

To begin to define the heterogeneity of BM-MSC populations, we combined CD271 with markers previously found to be expressed on human BM-MSCs. CD146 (MCAM) is a receptor for lamininα4 that is highly expressed in vascular muscle wall cells, endothelial cells and pericytes. CD271^+^CD146^+^ human MSC populations were found to have greater colony forming capacity compared to CD271^+^CD146^-^ populations^32^ and to be located in perivascular areas *in vivo*^19^. CD106 (VCAM-1) is a known adhesion molecule that facilitates interactions with vascular endothelium. We decided to combine these markers with VCAM-1 given the perivascular location of HSC niche cells in murine models, and because previous work showed that, within CD271+ BM MSCs, enrichment with VCAM-1 identifies self-renewing cells^21^. Importantly, single cell analysis of CD271+ MSCs has identified similar functional subsets within the MSC population as those we have identified here^33–36^. However, without the ability to identify these subsets immunophenotypically, it is impossible to directly test them functionally ex vivo. What we have demonstrated here through the ability to identify these cells by surface biomarker expression of CD271, CD146 and CD106, provides a novel tool for direct comparative analysis of these populations in human health and disease.

By combining transcriptional profiling with functional assays, we confirmed the heterogeneity of CD271^+^ BM-MSCs and identified CD271^+^CD146^+^CD106^+^, NVML cells, as the subset enriched for stem cell characteristics, as evidenced by a greater colony forming capacity, greater potential for multi-lineage mesenchymal differentiation and increased quiescence.

While there are considerable murine data regarding hematopoietic supportive MSC subsets, human MSC subsets have been difficult to test functionally for hematopoietic support. This is of critical importance, since emerging data strongly suggest that microenvironmental remodeling may contribute to clonal dynamics and evolution to leukemic transformation^37, 38^. Our work represents one of the first functional tests on hematopoietic supportive capacity of MSC subset populations.

Our data identify NVML cells as especially supportive of normal HSPCs (Figure 6). Consistent with this, in comparison to NLCs, the transcriptional profiles of NVML cells demonstrate greater overlap with the HSC-supportive murine LepR+ MSCs identified by Baryawno et al in a published murine BM single-cell RNAseq dataset^15^. Our analysis confirms that the BM stroma taxonomy identified in this murine studies is recapitulated by human MSC subpopulations. The fact that our newly defined human NVML cells are aligned with the well-characterized murine LepR+ MSC population represents a powerful tool to compare the impact of perturbations of the murine BMME to human samples in future studies.

**Figure 6.**
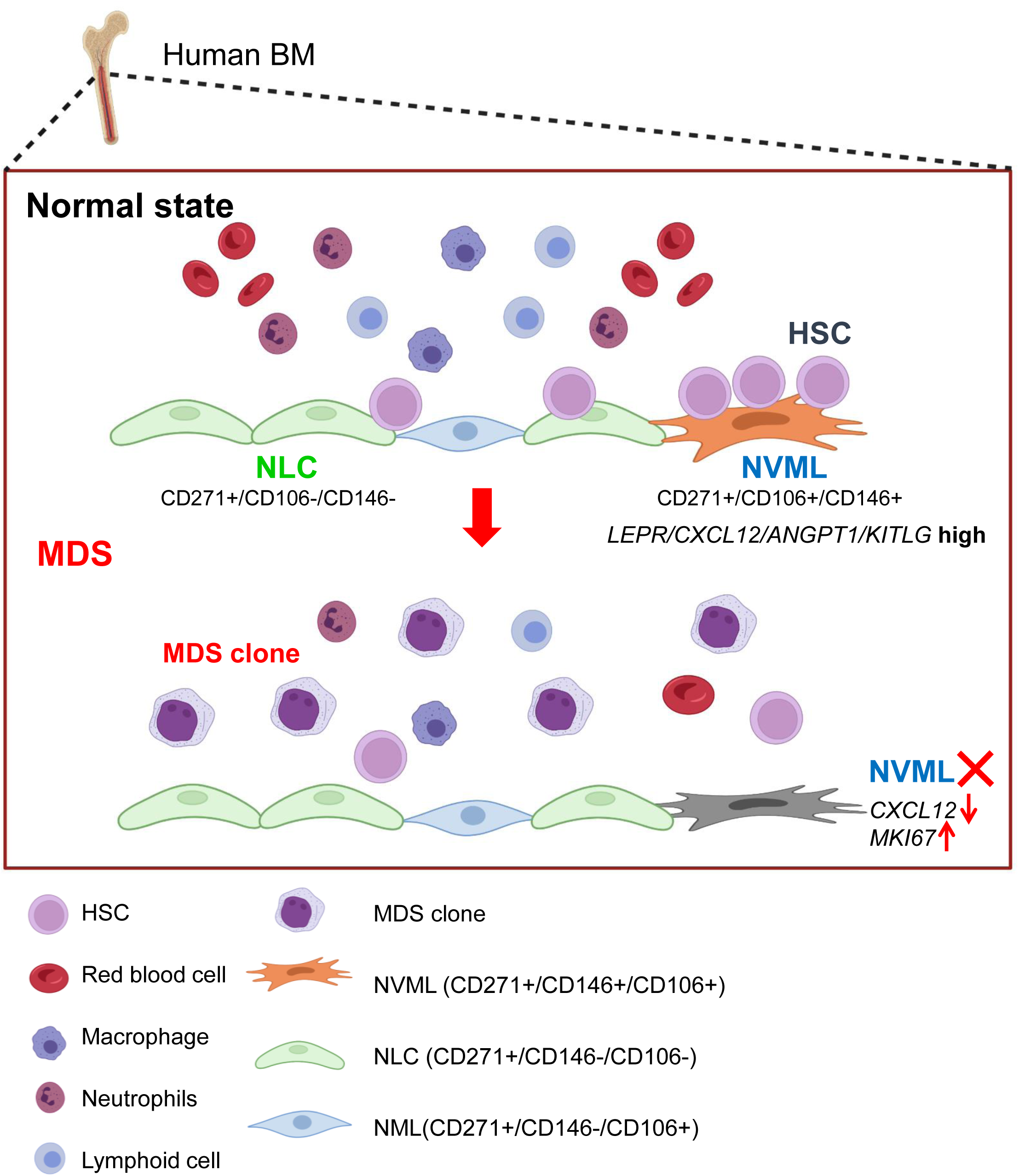
CD271, VCAM-1 and CD146 support normal hematopoietic stem and progenitor cells but are disrupted in myelodysplastic syndromes. Schematic summary of NVML niche cells in normal and MDS bone marrow.

Current strategies assessing functional MSC subsets ex vivo rely heavily on proliferative capacity and minor changes in trilineage differentiation capacity^33, 39, 40^. While these are characteristic functions of MSCs they do not address hematopoietic support. While there is considerable murine data regarding hematopoietic supportive MSC subpopulations^41, 42^, human subpopulations have been difficult to test functionally. To confirm the identity of human BM-MSCs capable of supporting HSPCs, we established co-culture systems of human NVML and NLC cells in the absence of supplemental cytokines. Notably, our sorting strategy using bone marrow aspirates and *ex vivo* expansion procedures made this functional assay possible. This approach is unique compared to previous studies that rely on getting transcriptional information from specific populations of MSCs after ex vivo expansion ^33, 34^ and *in vivo* histological evaluation^6^ or only utilized *ex vivo* culture using bone fragments rather than BM aspirates^21^. This innovation helps us highlight the supportive role of MSC populations that may at least in part be due to their production of growth factors and cytokines.

Analysis of MSC subsets in a diverse group of MDS patients demonstrates that the transcriptomes of MDS-derived NVML cells share patterns of dysregulated expression despite the mutational heterogeneity of the associated myeloid malignancy. Our transcriptional and functional data demonstrate that MDS-derived NVMLs lose their ability to support normal HSPCs, suggesting that disruption of NVML cells is at least partially responsible for the loss of normal hematopoietic function common to patients with MDS. We speculate that the loss of supportive niches for normal HSPC combined with the competitive advantage provided by the mutation(s) in the clones would result in acceleration of clonal expansion as a mechanism of evolution from clonal hematopoiesis to MDS and transformation to leukemia. Moreover, since one mechanism of treatment resistance is thought to be the emergence of novel clones, our ability to identify signals in NVML cells that are present in MDS patients with mutational heterogeneity may prove a superior therapeutic strategy that may reverse bone marrow failure while decreasing support of diverse MDS clones. Further studies are needed to define the mechanisms driving NVML dysfunction, however our data suggest that stem cell characteristics of NVML cells are critical for their ability to support HSPCs.

In summary, our study identifies a novel niche cell population in the human bone marrow that preferentially supports normal HSPCs and is disrupted in the setting of MDS (Figure 6), providing a new approach to dissect microenvironment-dependent mechanisms regulating homeostasis of human BMME with implications for clonal dynamics and progression to MDS.

## Acknowledgements

The authors would like to thank Lizz LaMere for technical assistance and the University of Rochester Flow Cytometry Core and Genomics Core for their assistance. Funding for this work was provided by the CTSI Incubator Program, a pilot from the UR Center from Rochester Resource-Based Center for Bone, Muscle and Orthopedic Research (ROCSTARR) to YK (P30AR069655), R01s AG079556 (to LMC and MWB) and AG076786 (to LMC), the Edward P. Evans Foundation (to LMC) and the Wilmot Cancer Institute.

## Author contributions

A. Y. Kawano, H.K., L.M.C., M.W.B., D.G designed the experiments, analyzed the data, interpreted the results and wrote the manuscript. Y. Kawano, H.K., M.A., M.W.L, T.C.H, and D.K.B. performed the experiments. D.T.S. Y.K., T.J.F. J.R.M., J.M.A. and J.L.L. contributed to data interpretation and discussion.

## Declaration of interests

The authors declare no competing interests.

**Additional information** is provided in Supplementary Materials.

## Availability of Data and Materials

The datasets used and/or analyzed during the current study are available from the corresponding authors.

## METHODS

### Patients

Seventeen patients with MDS (53 to 88 years old) were available for this study (supplemental Table 1). BM and peripheral blood samples were obtained at presentation to the University of Rochester Medical Center following informed consent under institutional review board-approved protocols. Bone marrow aspirates from eight healthy age-matched donors (Control BM, 5 males and 3 femals) were obtained after informed consent of an institutional review board-approved protocol as previously described^43^.

### Mononuclear cell isolation from bone marrow aspirates

Marrow aspirate material was handled as previously described^44^. Briefly, marrow was diluted in phosphate buffered saline (PBS, Gibco), layered with a Ficoll-Paque (GE Healthcare Bio-Sciences AB) gradient and centrifuged at 1,600 rpm at room temperature (RT) for 30 minutes. The light density bone marrow (LDBM) cells at the interface were collected and washed in 1X PBS and counted. Samples were cryopreserved with CS10 (STEMCELL technologies) in liquid nitrogen for further processing. Control BM Samples for the analysis of CFU-F and *ex vivo* expansion for differentiation capacity and co-culture (described below) were freshly processed.

### Cell sorting

For transcriptional analysis, cryopreserved LDBM cells were thawed and resuspended in FACS buffer (PBS with 2% FBS) and stained with biotinylated antibody cocktail (biotin-CD45/CD31/CD235ab) (1:50, 1:100, and 1:100 antibody dilution respectively, 200 μL per 1x10e6 cells) with human IgG-Fc block (1:100) followed by the incubation with streptavidin-magnetic bead particles (BD Biosciences, Ref#557812) at 50 μL per 1x10e6 cells. Magnetic particles were removed by using magnetic base based on the manufacturer’s direction. Depleted cells were stained with PerCP-Cy5.5-streptavidin, APC-Cy7-CD34, FITC-CD271, APC-CD106, PE-CD146, and DAPI. Human MSC subpopulations were sorted on a FACSAriaⅡ with 405-, 488-, 532- and 640-lasers (BD Biosciences) at the University of Rochester Medical Center Flow Core. Cells were sorted out for three populations as indicated in Figure 1A and submitted to RNA sequencing. For cell culture assays, LDBM cells were stained with APC-Cy7 antibody cocktail (CD45/CD31/CD34) and biotin-CD235ab that is subsequently stained with APC-Cy7-streptavidin. Cells labeled with APC-Cy7 were depleted by using anti-APC magnetic beads (Miltenyi Biotec, Ref#130-090-855) with auto MACS pro (Miltenyi Biotec). The selected APC- Cy7 negative population was subsequently stained with PE-CD271, PE-Cy7-CD146, and APC-CD106 and incubated at 4°C for 30 min. The gating strategy used to identify populations enriched for cells of interest is described in Result section as well (Figure 1A). Antibodies used in strategy are listed in Supplemental Table1.

### Colony forming unit of fibroblast (CFU-F) and cell culture

Sorted MSCs were collected in αMEM without ascorbic acid (Gibco, #A1049001) supplemented with 10% FBS, 1% P/S, and 25mM HEPES and plated for culture assay. The MSCs were plated at a minimal density of 500 cells/cm^2^, in αMEM without ascorbic acid supplemented with 10% FBS and 1% P/S on a Collagen-1 coated culture dish (Method described below). Cultures were incubated in hypoxic conditions (37^°^C, 2% O2, 5% CO2). Full media changes were performed every 3-4 days. Attached cells were passaged after approximately 80-90% confluence was attained. For passaging, cells were washed twice with 1X PBS, detached by incubating each well with TrypLE Express Enzyme (Gibco, #12605010) at 37°C for 4-5 minutes and split at appropriate cell densities. *Ex vivo* expanded MSCs were used at passage 3 to 6 in the subsequent experiments. For colony forming unit of fibroblasts (CFU-Fs) analysis, MSCs were plated (500 cells/cm^2^)^32^ in appropriate sized normal coated tissue culture plates and cultured in αMEM without ascorbic acid supplemented with 20% FBS and 1% P/S with a medium change on day 7 and day10 or day11. On day 14 the cells were fixed in 10 % neutral buffered formalin, stained with crystal violet and then colonies were defined by more than 40 cells/colony to be counted under the microscope.

### Flow cytometry

Analysis of marrow cell populations was performed as previously described^10^. Samples were run on a BD LSR Fortessa flow cytometer: 5 lasers, UV (355 nm), violet (405 nm), blue (488 nm), yellow-green (561 nm), and red (640 nm) lasers (BD Biosciences). As a viability stain, DAPI was used. Analysis was performed using FCS Express version 7 (De Novo Software).

### Plate coating with rat Collagen Type I

The tissue culture treated plates were coated with rat tail collagen Type I (Corning, Ref#354236) according to the manufacturer’s protocol. Briefly, 50 μg/ml of Collagen Type I solution was prepared using 0.02 N acetic acid and then dispensed as 5 μg/cm^2^ in the designated plates followed by overnight incubation at room temperature. On the following day, the plates were washed with PBS once and stored at 4°C.

### RNA-seq analysis

Total RNA was extracted from sorted MSCs using the RNeasy Plus Micro Kit (Qiagen, Valencia, CA) per manufacturer’s recommendations. RNA concentration was determined with the NanoDrop 1000 spectrophotometer (NanoDrop, Wilmington, DE) and RNA quality assessed with the Agilent Bioanalyzer 2100 (Agilent, Santa Clara, CA). 1ng of total RNA was pre-amplified with the SMARTer Ultra Low Input kit v4 (Clontech, Mountain View, CA) per manufacturer’s recommendations. The quantity and quality of the subsequent cDNA was determined using the Qubit Fluorometer (Life Technologies, Carlsbad, CA) and the Agilent Bioanalyzer 2100. 150pg of cDNA was used to generate Illumina compatible sequencing libraries with the NexteraXT library preparation kit (Illumina, San Diego, CA) per manufacturer’s protocols. The amplified libraries were hybridized to the Illumina flow cell using the cBot (Illumina, San Diego, CA) and sequenced with single-end reads of 100 nt on the Illumina HiSeq2500.

Raw sequencing files were demultiplexed using bcl2fastq-2.19.0. Quality filtering and adapter removal were performed using FastP-0.20.0. The resulting fastq files were mapped to the hg38 reference genome using STAR aligner version 2.7.0f. Gene-level read quantification was performed with featureCounts on exonic regions based on gencode M31 annotation. Differential expression analysis was performed using DESeq2-1.22.1 in R version 3.5.1. Differentially expressed genes were selected using a Wald test with adjusted p-value threshold of 0.05. Benjamini-Hochberg multiple test correction was applied to calculate the adjusted p-value. In the differential expression comparisons across all cell populations in normal bone marrow (NLC, NML and NVML), all Control samples were included in the regression model. NBM NML samples were excluded from the regression step in the NBM NLC and NVML cell population comparison. In the differential expression comparison of MDS versus Control samples, the NLC and NVML Control samples and the MDS samples were utilized in DESeq2 model building. All regression models were adjusted for sequencing batch. Principal component analysis was performed using the DESeq2 pcaplot function on rLog transformed expression values. Principal component plots were created using pcaExplorer-2.14.2 in R. Heatmaps were generated using pheatmap-1.0.12 on rLog transformed expression values and were scaled by expression value within each row. Gene set enrichment analyses was performed using the enrichR-3.0 R package. We tested for enrichment of both up and down regulated differentially expressed gene sets across terms in the wikiPathways human database version 2019, GO Biological processes version 2018, KEGG pathways human database version 2019, and the ChEA 2016 database. Genelists for Lepr+ MSC associated marker genes were obtained from scRNAseq data as previously published^15^.

### DNAseq variant analysis

Targeted sequencing analysis of 14 MDS patients BM samples was performed using Illumina TruSight Myeloid Sequencing Panel following the manufacturer’s recommendations. Libraries were sequenced on Illumina HiSeq2500 with PER250 to generate mean amplicon coverage depth of 15000X or greater. The threshold for quality control for any mutation identified was minimum of 20X read depth.

Samples were analyzed with the Illumina Basespace TruSeq amplicon workflow-3.0.0 using reference genome hg19. The workflow uses Isas-1.17.9.271+TSAv3, SAMtools-1.2, Isas Smith-Waterman-Gotoh-6.2.1.25 aligner, and Pisces-5.2.1.22 variant caller. Variants were annotated using the Illumina annotation engine-1.5.3.82 with annotation dataset-84.24.36. The readout for identified driver genes for MDS pathology were defined by in-house NGS method of University of Rochester listed in Supplemental Table 3B.

### MSC differentiation

MSCs were differentiated along osteogenic, adipogenic, and chondrogenic pathways using standard methodologies previously described^44^. Tri-lineage differentiation capacity assay was tested utilizing passage 4 to 6 cultures of sorted cells. Briefly, for the adipocytic induction, after MSCs reached 90-100% confluence, fat induction medium containing α-MEM supplemented with 10% FBS and 1% Penicillin-Streptomycin-glutamine (100X, Gibco), rosiglitazone (10µM; Cayman Chemicals), dexamethasone (1µM; Sigma-Aldrich), isobutyl-1-methylxanthine (IBMX; 25µM; Cayman Chemical) and insulin (0.2unit/ml; Humulin^®^ R; Lilly LLC) were added. Induction media were changed after three days and replaced with maintenance media containing insulin (0.2 unit/ml) for 24 hr. This cycle was repeated twice followed by 5 days of maintenance and at day12, cells were fixed and stained with Oil Red O solution (Electron Microscope Sciences). Images were taken with a light microscope (Olympus America) equipped with a camera (Olympus Inc.) and the percentage of MSCs positive for oil Red O was determined by manual counting of 3 randomized fields at 40X magnification from triplicated wells. For the osteogenic induction, after plated MSC cells reached 70-90% confluence, growth media were replaced with complete mineralizing media (α-MEM + ascorbic acid, supplemented with 10% FBS and 1% PSG, 10 nM dexamethasone (Sigma-Aldrich, St. Louis, MO, USA), 100 mM L-ascorbic acid (Sigma-Aldrich) and 10 mM β-glycerophosphate (Sigma-Aldrich). After 14 days of induction, cells were fixed and stained with 0.2% Alizarin Red S (Electron Microscope Sciences) to identify calcium deposition or stained for alkaline phosphatase activity and bond nodule formation following an alkaline phosphatase/Von Kossa staining. Chondrocyte induction was performed with MesenCult^TM^-ACF chondrogenic differentiation complete medium according to the manufacturer’s protocol (StemCell Technology). After 21 days of induction, cell pellets were fixed in 10% Neutral Buffered Formalin and embedded in Histogel (ThermoFisher Scientific). After embedding in paraffin and sectioning, Alcian-Blue (EMD Millipore) staining for cartilage was performed^45^ with nuclear staining of Nuclear Fast Red (R&D).

### Light microscopy

Images were taken at room temperature using CKX41 inverted microscope (Olympus) and DP74 camera (Olympus). Cellsens software (Olympus) was used to acquire images on the microscope.

### MSC co-culture to assess colony forming units (CFU-C)

CFU-Cs were assayed as previously described^46^. Cryopreserved LDBM cells from normal donors (Control BM: 1X10^5^ cells/well) or MDS-L cell line (1X10^3^ cells/well) were co-cultured in RPMI1640 supplemented with 10% FBS, 100U/ml PS, and 1x 2-mercaptoethanol (Gibco) with MSC monolayers which were 80%-90% confluent. All co-cultured cells were collected after trypsinization and then plated to duplicate dishes in MethoCult medium (Stem Cell Technologies, H4534) supplemented with 3 units/ml of human EPO (Peprotec, #100-64) and 10 ng/ml of G-CSF (Filgrastim, Millipore Sigma, 121181-53-1) (Stem Cell Technologies, Vancouver, BC, Canada).

After the 7-days for NBM and 14-days for MDS-L incubation at 37°C and 5% CO2, CFU-Cs or CFU-Es were defined by more than 40cells/colony (red-colored colonies were defined as CFU-Es in addition to that) to be counted manually under microscope.

### Statistical analysis

Data were expressed as mean ± standard error of the mean or ± range. Some analyses were made with GraphPad Prism software (version 9) using 2-tailed Student’s *t* test, Mann-Whitney nonparametric testing, or 1-way ANOVA with Tukey’s multiple-comparisons post-test or corrected method of Benjamini and Yekutieli when appropriate. A *P* value less than 0.05 was considered significant and denoted by asterisks (*p<0.05, **p<0.01, ***p<0.001). RNA sequencing data analysis were performed as described method in the Methods section (RNA sequencing), and adjusted p value(q) less than 0.05 was considered significant and denoted by asterisks (*q<0.05, **q<0.01, ***q<0.001). Statistical methods are also described in figure legends.

**Supplemental Figure.1**

A. Plot of first (PC1) and second (PC2) principal components from principal component analysis of RNA expression in NBM samples. NLCs (green), and NVML cells (blue) samples and NML cells (orange) are shown. B. Hierarchical clustering heat map of differentially expressed genes in NVML cells vs NML cells vs NLCs samples scaled per gene. n=8

**Supplemental Figure.2**

Heatmap of marker genes for LepR+ MSCs in scRNAseq murine dataset^15^ including all three populations NLCs, NML cells, NVML cells.

**Supplemental Figure 3.**

Heat map of variant allele frequency (VAF) of mutated genes for MDS patients’ BM samples.

**Supplemental Figure 4.**

Transcriptional impact of MDS on MSC subsets A. Upregulated and B. downregulated gene enrichment results from differentially expressed genes in MDS NVML cells vs Control NVML cells. Only the top 30 (A) and 11 (B) hits are shown. Circle size represents higher proportion of differentially expressed genes enriched for that term. C. Hierarchical clustering heat map created by GSEA in Control (C_) NLCs (Green) vs Control (C_) NVML cells (Orange) vs MDS NVML cells (light blue) samples scaled per gene of Cellular senescence and Fanconi anemia pathways. D. Normalized counts of transcriptomes upregulated in MDS samples in MDS NVML cells vs Control NVML cells. E. Normalized counts of downregulated transcriptomes in MDS samples in Control (Cont) NVML cells vs MDS NVML cells. n=8-17, Min to max, adjusted p(q) or p value from Wald test (p) are provided.

**Supplemental Table.1**

List of the antibodies used in experimental procedures.

**Supplemental Table.2**

Clinical and cytogenetic characteristics of MDS patients in the samples sorted for RNAseq analysis of MSCs.

**Supplemental Table.3**

Molecular mutations detected in bone marrow of MDS patients.

**Supplemental Table.4**

Tables of defined driver genes for MDS pathology performed by in-house NGS method of University of Rochester.

