## Supplemental Figures for "Myelodysplastic syndromes disable support of human hematopoietic stem and progenitor cells by CD271, VCAM- 1 and CD146 expressing marrow niche cells"

### Supplemental Figure.1

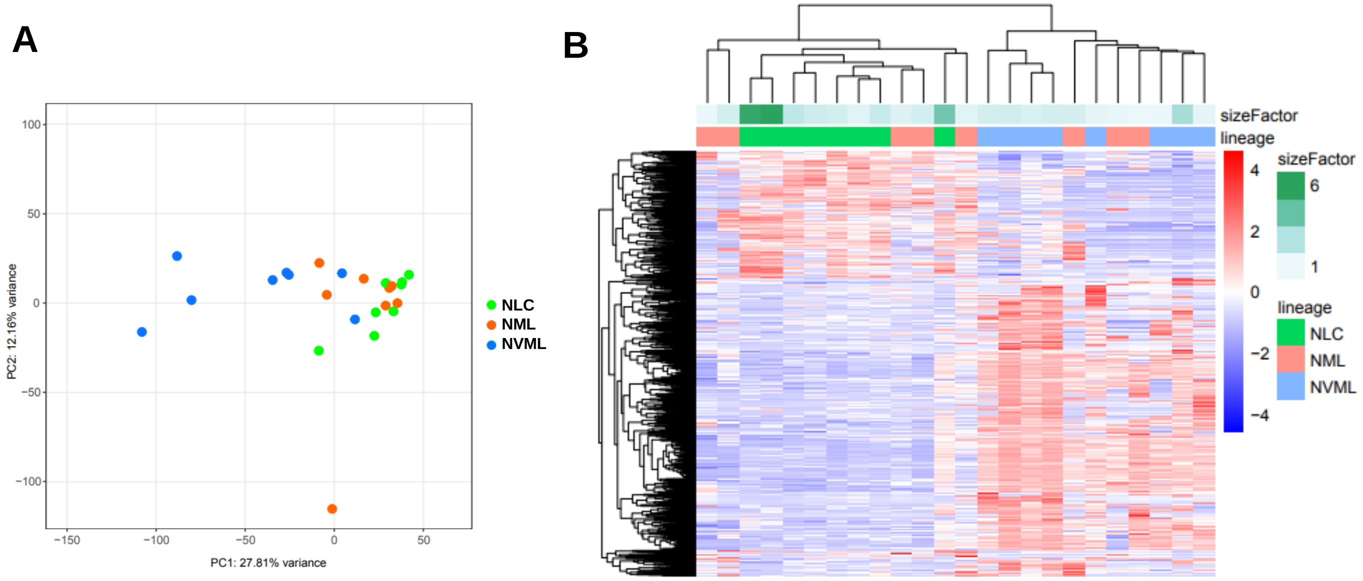

### Supplemental Figure.2

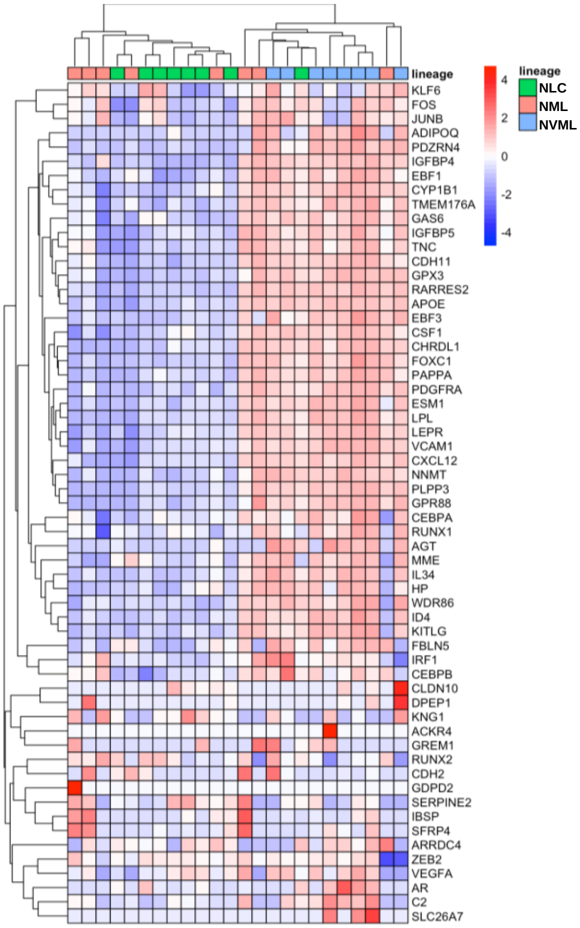

#### Supplemental Figure.3

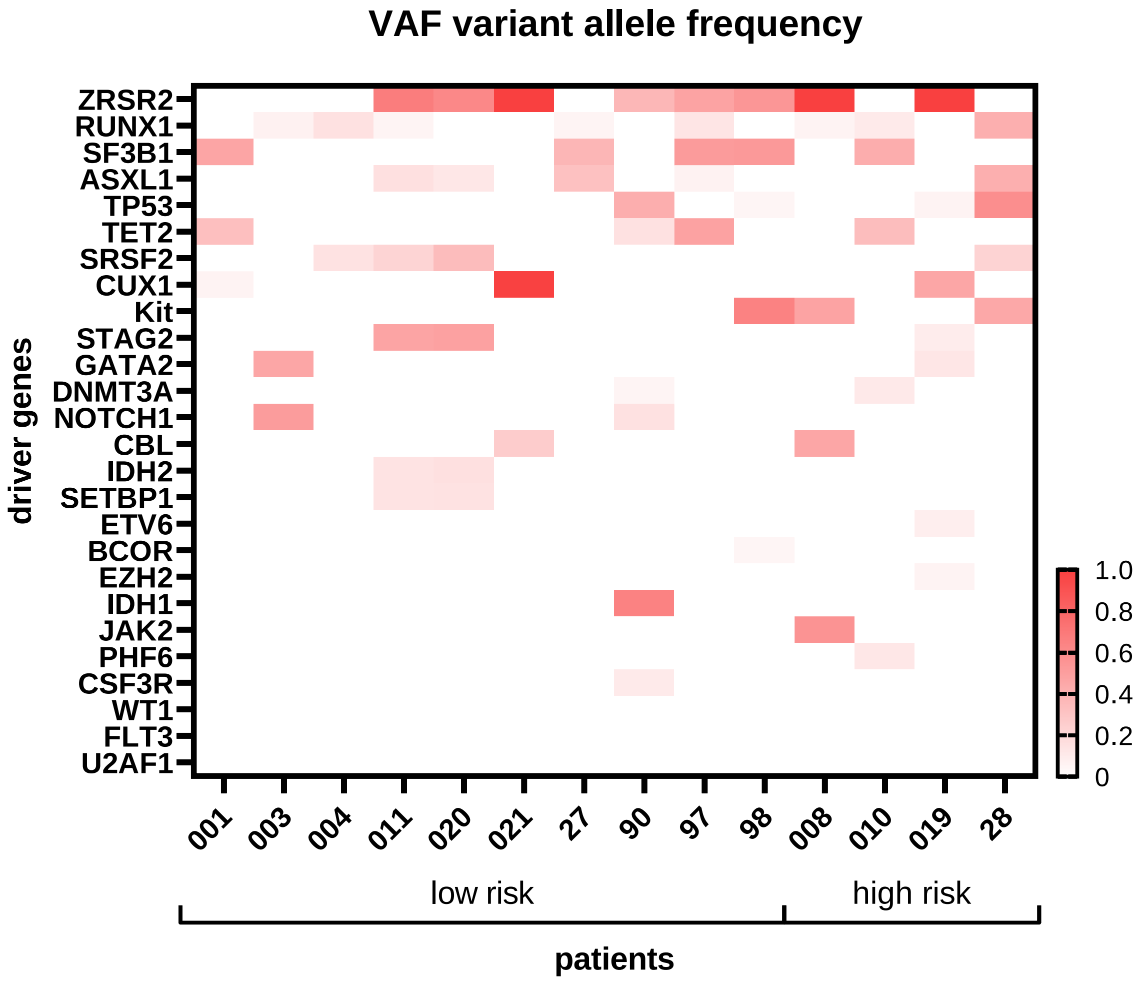

### Supplemental Figure.4

**A** MDS NVML vs NBM NVML up-regulated pathway

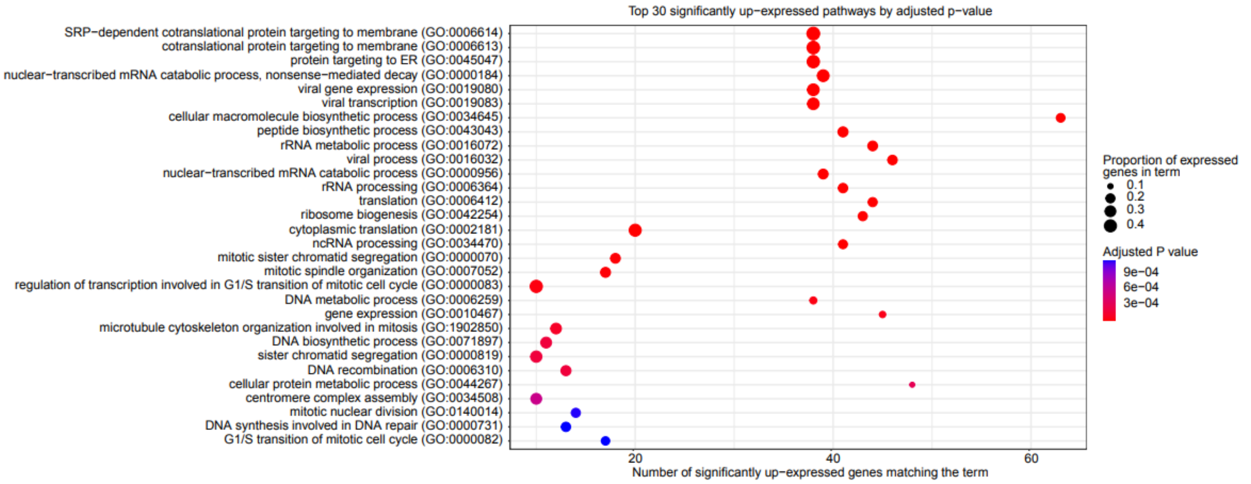


**B** MDS NVML vs NBM NVML down-regulated pathway

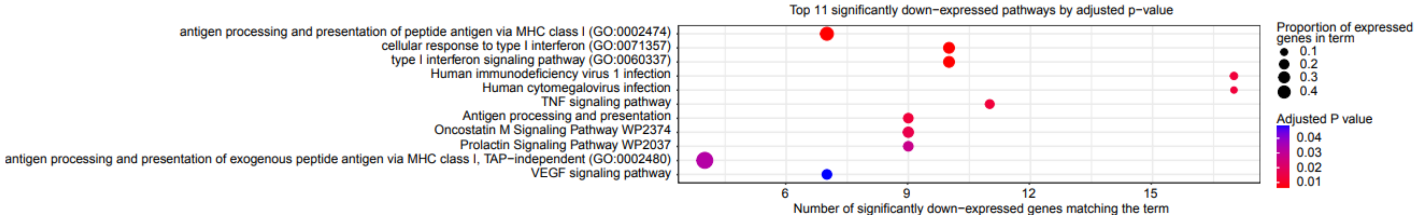


**C**

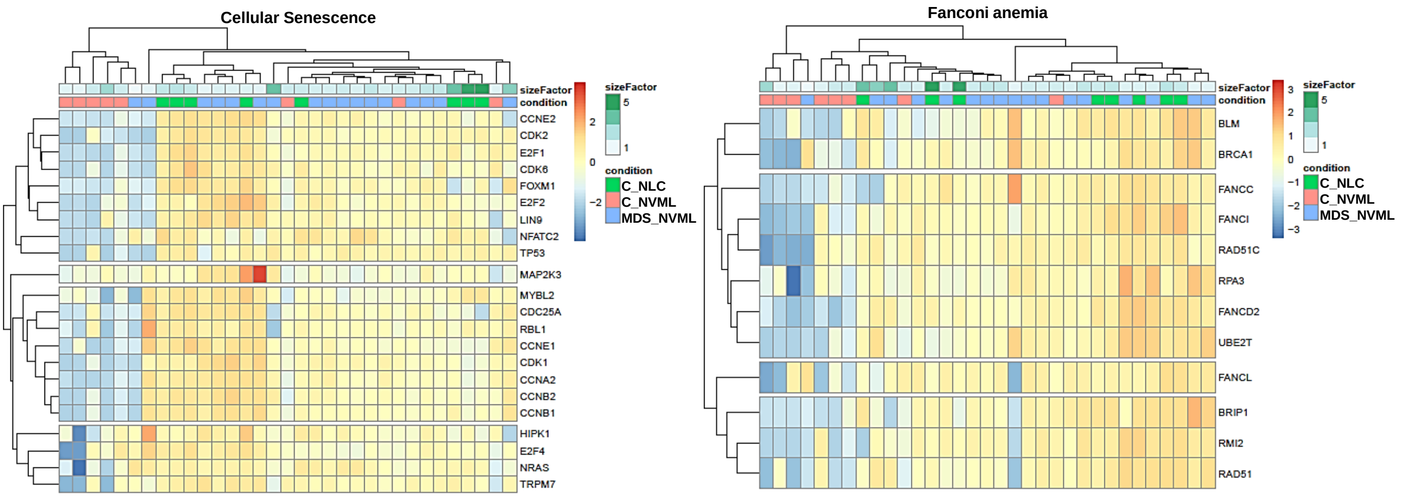


**D**

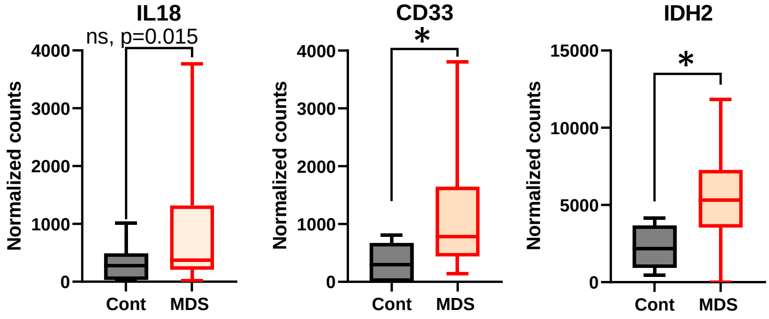


**E**

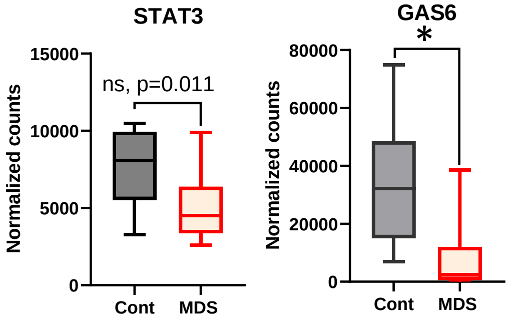
