## supplemental table1 for "Myelodysplastic syndromes disable support of human hematopoietic stem and progenitor cells by CD271, VCAM- 1 and CD146 expressing marrow niche cells"

**Supplemental Table.1**

| **Antibody** | **Company** | **Ref#** | **Clone** |
| --- | --- | --- | --- |
| biotin-antiCD45 | eBioscience | 13-0459-82 | HI30 |
| biotin-antiCD31 | eBioscience | 13-0319-82 | WM59 |
| biotin-antiCD235ab | eBioscience | 13-9987-82 | HIR2 |
| PerCP-Cy5.5-streptavidin | BD Pharmingen | 551419 |  |
| FITC-antiCD271 | BioLegend | 345104 | ME20.4 |
| APC-antiCD106 | BioLegend | 305810 | STA |
| APC-Cy7-antiCD34 | BioLegend | 343514 | 581 |
| APC-Cy7-antiCD45 | BioLegend | 304014 | HI30 |
| APC-Cy7-antiCD31 | BioLegend | 303120 | WM59 |
| APC-Cy7-streptavidin | BD Pharmingen | 554063 |  |
| PE-antiCD146 | BD Pharmingen | 561013 | P1H12 |
| PE-antiCD271 | BioLegend | 345106 | ME20.4 |
| PE-Cy7-antiCD146 | BioLegend | 361008 | P1H12 |
| DAPI | Invitrogen | D21490 |  |

**Supplemental Table.2**

| **MDS ID** | **Gender** | **Age** | **Risk** | **Diagnosis** | **Karyotype** |
| --- | --- | --- | --- | --- | --- |
| MDS001 | M | 71 | Low | MDS-RS-MLD | 47, XY, +8 |
| MDS003 | M | 73 | Low | MDS-MLD | 46, XY |
| MDS004 | M | 79 | Low | CMML-1 | 46, XY |
| MDS005 | M | 57 | High | MDS-EB-1 | 46, XY |
| MDS008 | M | 77 | High | MDS-EB-2 | Complex |
| MDS010 | M | 53 | High | MDS-EB-1 | 46, XY |
| MDS011 | M | 75 | Low | MDS-MLD | Complex |
| MDS019 | M | 71 | High | MDS-EB-2 | 46, XY |
| MDS020 | M | 86 | Low | MDS-MLD | 46, XY,i(17)(q10) |
| MDS021 | M | 69 | Low | MDS-MLD | 45, X, -Y |
| MDS027 | M | 62 | Low | MDS-MLD | 46,XY,del(11)(q23.2) |
| MDS028 | M | 75 | High | MDS-EB-2 | Complex |
| MDS031 | F | 65 | High | MDS-EB-1 | 46, XX, del(5)(q22q35) |
| MDS056 | M | 72 | Low | MDS-MLD | 46, XY, del(20)(q11.2) |
| MDS090 | F | 69 | Low | MDS-MLD | Complex |
| MDS097 | F | 72 | Low | MDS-RS-MLD | 46, XX |
| MDS098 | F | 88 | Low | MDS-RS-MLD | 47, XX, +8 |

**Supplemental Table.3**

| **MDS ID** | **Mutation** |
| --- | --- |
| MDS001 | SF3B1, TET2, CUX1 |
| MDS003 | RUNX1, GATA2, NOTCH1 |
| MDS004 | SRSF2, RUNX1, |
| MDS005 | N/A |
| MDS008 | ZRZS2, KIT, JAK2, CBL, RUNX1 |
| MDS010 | DNMT3A, SF3B1, TET2, RUNX1, PHF6 |
| MDS011 | IDH2, SRSF2, SETBP1, ASXL1, RUNX1, STAG2 |
| MDS019 | GATA2, CUX1, EZH2, ETV6, TP53, ZRSR2, STAG2 |
| MDS020 | IDH2, SRSF2, SETBP1, ASXL1, ZRSR2, STAG2 |
| MDS021 | CUX1, CBL, ZRSR2 |
| MDS027 | SF3B1, ASXL1, RUNX1 |
| MDS028 | KIT, TP53,SRSF2, ASXL1, RUNX1 |
| MDS031 | (JAK2V617F)# other mutations not assessed |
| MDS056 | N/A |
| MDS090 | CSF3R, DNMT3A, TET2, TP53, ZRSR2, IDH1, NOTCH1 |
| MDS097 | SF3B1, TET2, ASXL1, RUNX1, ZRSR2 |
| MDS098 | SF3B1, KIT, TP53, ZRSR2, BCOR |

**Supplemental Table.4**

| **Gene** | **Accession#** | **Exon(s)** |
| --- | --- | --- |
| ASXL1 | NM_015338.5 | 12 |
| CBL | NM_005188.3 | 8,9 |
| ETV6 | NM_001987.4 | all |
| FLT3 | NM_004119.2 | 14,15,20 |
| IDH1 | NM_005896.2 | 4 |
| KIT | NM_000222.2 | 2,8-11,13,17 |
| MYD88 | NM_002468.4 | 3-5 |
| NRAS | NM_002524.4 | 2,3 |
| RUNX1 | NM_001754.4 | all |
| SRSF2 | NM_00195427.1 | 1 |
| TP53 | NM_000546.5 | 1-11 |
| ZRSR2 | NM_005089.3 | all |

| **Gene** | **Accession#** | **Exon(s)** |
| --- | --- | --- |
| BCOR | NM_001123385.1 | all |
| CSF3R | NM_156039.3 | 14-17 |
| EZH2 | NM_004456.4 | all |
| GATA1 | NM_002049.3 | 2 |
| IDH2 | NM_002168.2 | 4 |
| KRAS | NM_033360.2 | 2,3 |
| NOTCH1 | NM_017617.3 | 26-28,34 |
| PHF6 | NM_032458.2 | all |
| SETBP1 | NM_015559.2 | 4 |
| STAG2 | NM_001042749.1 | all |
| U2AF1 | NM_001025203.1 | 2,6 |

| **Gene** | **Accession#** | **Exon(s)** |
| --- | --- | --- |
| BRAF | NM_004333.4 | 15 |
| DNMT3a | NM_022552.4 | all |
| FBXW7 | NM_033632.3 | 9-11 |
| GATA2 | NM_032638.4 | 2-6 |
| JAX2 | NM_004972.3 | 12,14 |
| MPL | NM_005373.2 | 10 |
| NPM1 | NM_002520.6 | 12 |
| PTPN11 | NM_002834.3 | 3,13 |
| SF3B1 | NM_012433.2 | 13-16 |
| TET2 | NM_001127208.2 | 3-11 |
| WT1 | NM_024426.4 | 7,9 |
